# Development of immunological reagents for the identification of biomarkers of inflammation in the common marmoset

**DOI:** 10.64898/2026.09.17.752376

**Authors:** Ektoras Lambrou, David Hallengärd, Benjamin Turbow, Jessica Callery, Vida Hodara, Jonathan King, Corinna Ross, Niklas Ahlborg, Bartek Makower, Luis D. Giavedoni

## Abstract

Marmosets (MAR, Callithrix jacchus) are nonhuman primates extensively studied in a broad array of preclinical biomedical research areas. However, the limited number of available immunological reagents remains a critical shortcoming of these models. We successfully developed monoclonal antibodies (mAbs) pairs that can be used in immunoassays for the identification and quantification of the biomarkers of inflammation C-reactive protein (CRP), CXCL-10 (IP-10), interleukin (IL)-4, IL-6, and IL-10. Recombinant MAR protein variants were produced in mammalian cells, purified and used for mice immunization. Hybridoma clones were selected first for their capacity to bind the recombinant proteins. All possible pair combinations of these mAbs were then tested for their capacity to identify their targets in MAR-derived samples. These MAR samples consisted of supernatant of PBMCs exposed ex vivo to different stimulants, and plasma samples from animals stimulated in vivo with a low-dose intravenous LPS inoculation. The majority of the mAbs that recognized the recombinant proteins failed to bind to the MAR-derived homologs. However, we found MAR-derived reactive mAbs and selected pairs with optimal sensitivity and signal-to-noise ratio for immunoassays that included Luminex multiplexing assays, ELISA, and ELISpot assays. These immunoreagents and assays will improve the translational value of different MAR biomedical models.

## Introduction

Non-human primate (NHP) models have been extremely useful in scientific research due to their similarity to the immune system of humans and susceptibility to infection with many human pathogens (Tarantal et al., 2022, Roozendaal et al., 2020, Phillips et al., 2014, Ishigaki and Itoh, 2025, Berry et al., 2024). As a result, NHPs constitute valuable animal models for studying infectious diseases; social, cognitive and behavioral research; reproductive biology; regenerative medicine; aging and neuroscience, and preclinical evaluation of new therapeutics as well as for assessment of vaccine-induced immunity. Marmosets (MAR, Callithrix jacchus) are small South American NHPs studied extensively in a broad array of research areas, including infection, vision, audition, social behavior, cognition, neurodegenerative diseases, reproduction, and genetic manipulation (Hosoya, 2025, Chen et al., 2025, Reveles et al., 2024, Peters et al., 2023, Perez-Cruz and Rodriguez-Callejas, 2023, Inoue et al., 2023, Singh et al., 2021, Tardif, 2019, Herron et al., 2024). One critical shortcoming of MAR biomedical models is the limited number of immunological reagents and immunoassays that are currently available for several biomarkers of inflammation. It is estimated that while the human lineage diverged from African and Asian NHPs about 23 million years ago (MYA; range 21-25), this separation is estimated to be 33 MYA (range 32–36) for American NHP (Glazko and Nei, 2003). Thus, due to this divergence, the amino acid differences between some human and MAR proteins make many monoclonal antibodies (mAbs), developed for the identification of human biomarkers, unable to bind to the same epitopes in MAR homolog proteins.

During this investigation, we produced recombinant variants of the MAR biomarkers of inflammation C-reactive protein (CRP), CXCL10 (C-X-C motif chemokine ligand 10, also known as interferon gamma-induced protein 10, IP-10), interleukin (IL)-4, IL-6, and IL-10. We used some of these variants to develop murine monoclonal antibodies (mAbs), which were first selected using recombinant forms of the MAR proteins. We finally tested different mAb pairs for their capacity to identify the natural form of these MAR proteins, and developed immunoassays to determine the presence of these biomarkers in MAR samples generated ex vivo and in vivo.

## Materials and Methods

### Eukaryotic expression of recombinant MAR CRP, CXCL-10 (IP-10), IL-4, IL-6, and IL-10

Recombinant MAR molecules were produced with overlapping oligonucleotides coding for the amino acid sequences obtained from Uniprot (http://www.uniprot.org) for CRP (F7IB03), CXCL-10 (A0A2R8MWC4), IL-4 (A4ZXA4), IL-6 (Q0Z973), and IL-10 (Q0Z972) as previously described (Hoglind et al., 2017). The MAR proteins were expressed as different variants, depending on their desired application (**Supplemental Table 1**). For immunizations, proteins were expressed in their native form (V1) and/or as fusion proteins designed to enhance immunogenicity in mice: a fusion to a proprietary cassette of promiscuous, H-2d-restricted T-helper epitopes (variant V4), or a fusion to one of two proprietary xenogeneic carrier proteins (variants V5a and V5b) (Mabtech AB, Nacka Strand, Sweden). Fusion of promiscuous T-helper epitope cassettes, or of heterologous immunostimulatory proteins, to weak immunogenic antigens is an established strategy for enhancing antigen-specific antibody responses (Alexander et al., 1994, Hung et al., 2007, McCormick et al., 2001, Nimal et al., 2005). The identities and sequences of the T-helper epitope cassette and of the xenogeneic carrier proteins are proprietary to Mabtech AB and are not disclosed; they are not required for the interpretation of the antibody characterization data presented here. The complete set of recombinant antigen variants, the codes used to designate them, the tags they carry, and their intended use are summarized in **Supplemental Table 1**, and the corresponding expression plasmids are listed in **Supplemental Table 2**. The variant codes defined in **Supplemental Table 1** are used consistently throughout the text, figures and tables. For IL-4 and IL-6, cytokines known to interact with soluble forms of their receptors, we also produced receptor silent variants, designated as ΔN. The corresponding soluble receptor ectodomains were expressed as human IgG4 Fc fusions and are designated MAR sIL-4Rα-Fc and MAR sIL-6Rα-Fc. All proteins were expressed in HEK-293 mammalian cells to ensure high similarity with endogenous MAR proteins.

### Production of monoclonal antibodies to MAR CRP, CXCL-10, IL-4, IL-6, and IL-10

MAbs to MAR CRP, CXCL-10 (IP-10), IL-4, IL-6, and IL-10 were induced in mice using methods previously described (Zuber et al., 2005). The mice were housed at the Karolinska Institute (Solna, Sweden) and handled in accordance with the guidelines of the Swedish Ethical Committee for Animal protection. Briefly, BALB/c mice were immunized with purified, recombinant HEK-derived MAR biomarkers in Immune-Stimulating Complex (ISCOM)-based adjuvant (NanoQuil, CRODA, Vejle, Denmark). Mice with the highest anti-MAR biomarker antibody responses received a booster immunization with the same antigen dose intraperitoneally without adjuvant. Spleens from these mice were collected three days later to prepare splenocytes for subsequent fusion with the mouse Sp2/0-Ag14 myeloma cell line. Subsequent screening was done by ELISA.

### Primary screening ELISA

Hybridoma supernatants were analyzed by ELISA for antigen reactivity. For this purpose, 96-well polystyrene plates were coated overnight with 1 μg/ml goat-anti mouse IgG in PBS and then blocked with PBS containing 0.1% bovine serum albumin (BSA, incubation buffer -IB-). Hybridoma supernatants were diluted in IB for 2 hs at room temperature, followed by Twin Strep-tagged antigen titrated in IB. After washing, Strep-tactin-HRP was added at 1:4000 dilution in IB, incubated for 1h, and antigen-antibody complexes were developed with TMB for 15 min at RT, followed by 0.18M SO_4_H_2_.

### Epitope mapping of mAbs

Positive hybridomas with high affinity were subcloned and mAbs were purified from supernatants on Protein G columns. A part of the purified mAb was biotinylated. All the selected mAbs were further characterized for monoclonality and IgG subclass by FluoroSpot assay (Hoglind et al., 2017). Epitope binning was performed using BioLayer Interferometry (BLI, OctetRED96e). BLI measures association/dissociation between mAbs and target antigen in real time (Abdiche et al., 2014).

### Ex vivo production of marmoset-derived samples

Blood samples (2 ml in EDTA) were collected from 40 adult MAR (20 from each sex) housed at the Southwest National Primate Research Center (SNPRC), Texas Biomedical Research Institute, San Antonio, TX, USA. Peripheral blood mononuclear cells (PBMC) were purified by centrifugation over Ficoll-Hypaque and stimulated as described before (Giavedoni, 2005). Briefly, PBMC were resuspended at 2×10^6^ cells/ml and maintained in RPMI 1640 supplemented with 10% fetal calf serum (RPMI-10). Cells were stimulated with a combination of phorbol myristate acetate (PMA, 50 ng/ml) and ionomycin (1 μg/ml), lipopolysaccharide (LPS, 0.5 μg/ml), phytohemagglutinin (PHA, 5 μg/ml), or Staphylococcus enterotoxin B (SEB, 1 μg/ml). Cell-free culture supernatants were harvested at 24 hours of stimulation, divided into aliquots and stored at -80°C.

### In vivo production of marmoset-derived samples

LPS injection has been used in MAR before as a way of inducing an acute inflammatory response (Gengozian et al., 1978, Philippens et al., 2017). Six adult animals (3 males and 3 females), housed at the SNPRC and following procedures approved by the Texas Biomed Institutional Animal Care and Use Committee (IACUC), received an intravenous injection of a sterile solution of Escherichia coli 0111:B4 LPS (LPS-EB VacciGrade, InvivoGen, San Diego, CA) in saline, in a concentration of 10 ng/kg of body weight (a dose previously applied to humans (Kaneko et al., 2003)). Approximately 150 μl of blood was collected in EDTA tubes at time-points 0 min, 30 min, 1 hour, 2 hours, 4 hours, 6 hours, and 24 hours. Plasma was separated by centrifugation, aliquoted and stored -80°C.

### ELISA with MAR-derived samples

To identify working mAbs pairs, we coated ELISA plates with a single capture mAb and then tested different mAbs against the same samples. Recombinant HUM cytokine standards (Mabtech), HEK-derived supernatants containing recombinant MAR cytokines, or stimulated MAR PBMC supernatants were added and incubated for 2 h at RT. The same sample pattern was repeated as many times as different detection mAbs needed to be tested. Next, biotinylated mAbs against MAR cytokines were added and incubated for 1 h at RT. Following that, streptavidin-horse radish peroxidase conjugate (SA-HRP; Mabtech) diluted 1:1000 in incubation buffer was added (100 µl/well) and incubated for 1 h at RT. The assay was developed with TMB substrate (Mabtech) and stopped with H_2_SO_4_ followed by absorbance measurement at 450 nm with an ELISA reader (SpectraMax, Molecular Devices, San Jose, CA, USA).

### ELISpot assay for MAR IL-4

ELISpot was performed essentially as described (Dillenbeck et al., 2014) using ethanol-activated polyvinylidene fluoride 96-well plates (Millipore Sigma, Burlington, MA. USA) coated with 100 μl of mAbs to MAR IL-4 at 15 μg/ml. MAR PBMC at 5 x 10^4^ cells/well were added in 100 μl of RPMI-10 with or without PMA (50 ng/ml) and ionomycin (1 μg/ml), or PHA (5 μg/ml). PBMC were incubated at 37 °C and 5% CO_2_ in humidified air for 48 h. After incubation, the plates were washed in PBS and incubated with biotinylated detection mAbs to MAR IL-4 at room temperature (RT) for 2 h, followed by incubation with streptavidin alkaline phosphatase (SA-AP) and subsequently NBT/BCIP substrate (Mabtech). ELISpot assays were set up in duplicates. An ELISPOT reader (Cellular Technology Ltd., Shaker Heights, OH, USA) was used to enumerate the spots representing cytokine-producing cells.

### Luminex assay for MAR cytokine/chemokine analyses

Cytokine concentrations in MAR plasma and PBMC supernatant were assessed by using the Luminex system as described previously for marmosets and other nonhuman primates (Mustoe et al., 2026, Peters et al., 2023, Ross et al., 2019, Giavedoni, 2005). The Luminex assay included evaluation of the following 5 MAR analytes: interferon gamma (IFN-γ), interleukin-6 (IL-6), IL-10, interferon-induced protein 10 (IP-10, CXCL10), and tumor necrosis factor-alpha (TNF-α). Previously established mAbs to human (HUM) IFN-γ (MT126L and 7-B6-1, MabTech) and TNF-α (MT21A8 and MT15B15, MabTech), known to cross-react with MAR cytokines, were added to the multiplex cocktail. The C-tag forms of MAR IL-6 and IP-10, as well as recombinant human IFN-γ, IL-10, and TNF-α molecules (Mabtech) were used as standards.

## Results

### Expression of recombinant MAR proteins

We expressed different variants of MAR biomarkers of inflammation in HEK cells. For immunizations, proteins were expressed in their native form (V1) and/or as immunogenicity-enhanced fusion variants (V4, V5a, V5b, and V6; **Supplemental Table 1**). C-tag or Fc domains were included to facilitate production. For serology and identification of positive hybridomas, the proteins were expressed with a Twin-Strep-tag at either terminus. Twin-Strep-tagged variants are designated V3N and V3C. In general, levels of expression varied for the different molecules and, for each cytokine, with the type of modification that was used (**Figure 1**). For example, the best expressing form of MAR IL-10 was a protein fusion with the human IgG_4_ Fc region (V6), which was purified with Protein G column (**Supplemental Figure 1**) and used for immunization. Expression levels for variants of MAR CRP, IL-4, and IL-6 were acceptable, but for MAR IP-10 only the V5a variant was produced.

**Figure 1.**
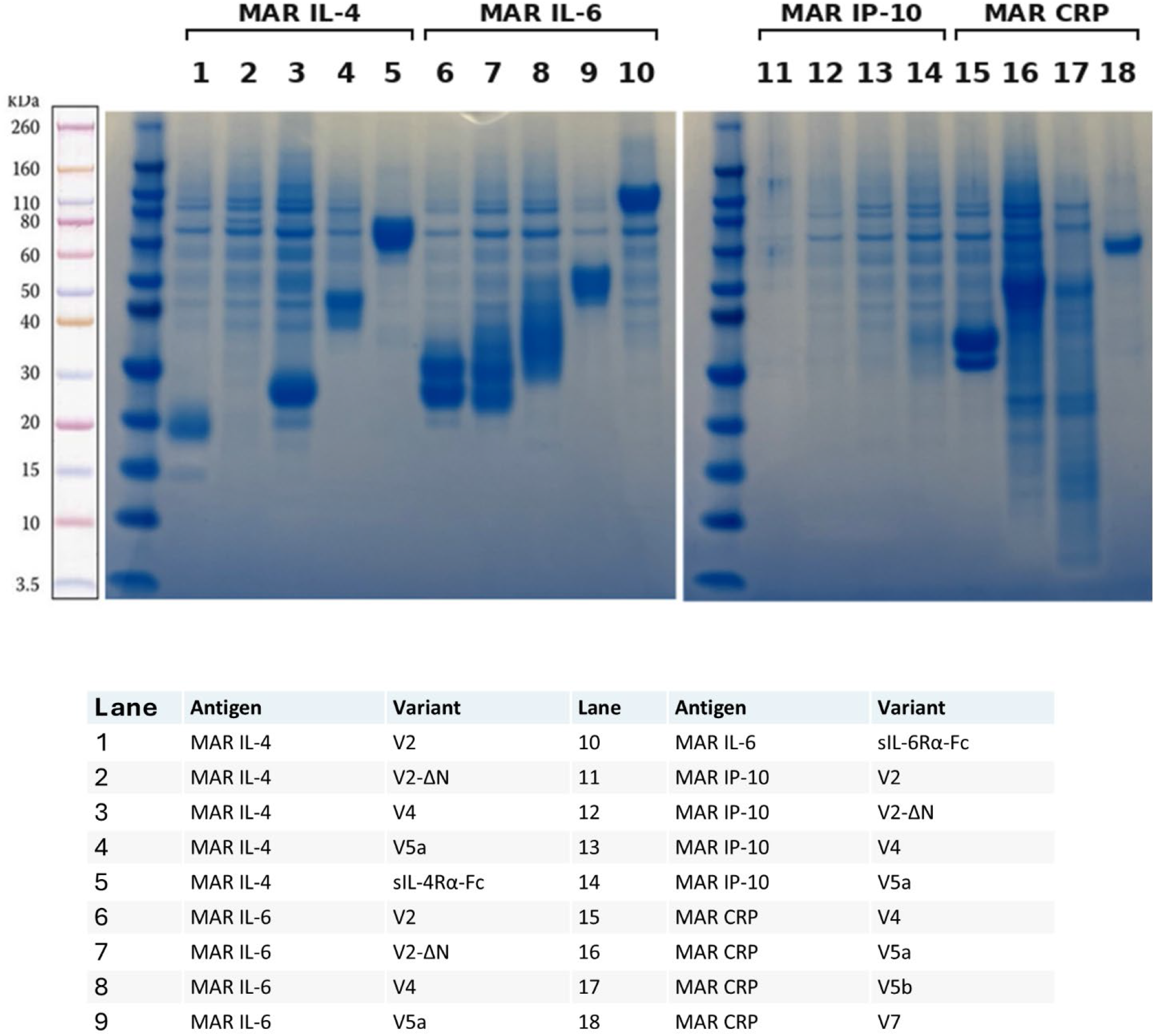
Expression of recombinant MAR antigen variants. SDS-PAGE with Coomassie blue staining of affinity-purified recombinant MAR IL-4, IL-6, IP-10 and CRP variants, and of the soluble IL-4 and IL-6 receptor ectodomains, all expressed in HEK-293 cells. Lanes are numbered and identified in the key; the variant codes are defined in Supplemental Table 1 and are used consistently throughout the manuscript. Variants V4, V5a, V5b and V7 carry proprietary immunogenicity-enhancing elements of Mabtech AB whose identity and sequence are not disclosed (see Author Disclosure Statement). Calculated molecular weights and purity for each variant are given in Supplementary Table 2.

### Generation of anti-MAR mAbs and epitope mapping

The different MAR proteins generated for immunization were administered to mice with an ISCOM-based adjuvant to generate anti-MAR biomarker mAbs. Following four immunizations, serology was performed using ELISA utilizing the Twin-tagged versions (N-terminal and C-terminal versions) of each biomarker. The reactive mAb clones and the epitopes they recognized were analyzed by biolayer interferometry using MAR molecules (**Figure 2**). For MAR CRP, 30 reactive clones were initially selected, and they were assigned to 9 different epitope groups; later, based on hybridoma productivity, a subset of 15 mAbs were grouped into seven epitopes. For MAR IP-10, 30 reactive clones were initially selected, and then a subset of 19 mAbs were assigned to six different epitopes. For MAR IL-4, 30 reactive clones were reduced to 10 mAbs that mapped to four different epitopes; an additional anti-human IL-4 mAb (IL-4I), recognized a 5^th^ epitope in MAR IL-4. For MAR IL-6, 40 clones were first selected, and nine mAbs mapping to five different epitopes were further evaluated. Finally, for MAR IL-10, eight hybridomas were identified and five were further characterized and found to bind to three different epitopes. All the selected mAbs were further tested in immunoassays with MAR-derived samples.

**Figure 2.**
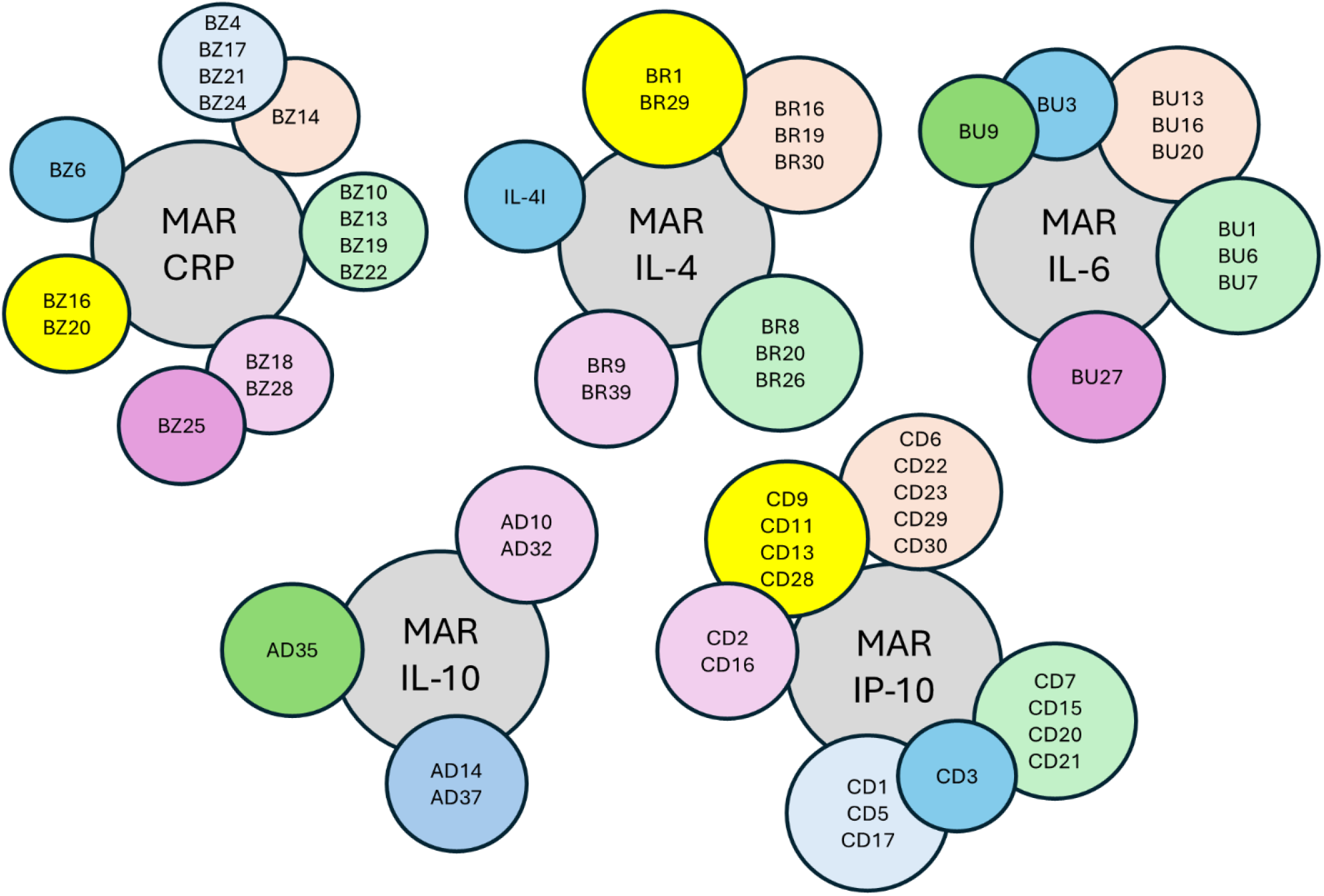
Epitope mapping for the mAbs produced against MAR biomarkers of inflammation. Epitope mapping was performed by BioLayer Interferometry using antibody pairs and soluble MAR biomarkers without tags. Each circle represents a different epitope containing all the mAbs targeting that epitope.

### Screening of selected mAbs with MAR-derived samples

MAR CRP: The result of the screening ELISAs indicated that all the 13 selected mAbs recognized the recombinant MAR CRP molecule. However, only a subset of mAb combinations recognized the native CRP molecule present in the plasma of marmosets challenged with LPS (**Figure 3A**). Further testing of reactive mAb pairs included MAR-derived samples as well as archived rhesus macaque plasma samples and recombinant human CRP (**Figure 3B**). We observed that different mAb pairs had different capacity to bind to proteins of different origin. For example, the mAb antibody pair BZ22/BZ24 (capture/biotinylated detection antibodies) detected all the tested samples, including the human and marmoset CRP standards, and CRP in the plasma of marmoset and rhesus macaques. Meanwhile, BZ13/BZ24 and BZ19/BZ25 only recognized recombinant and natural MAR CRP, while the pairs BZ4/BZ19, BZ10/BZ19, BZ19/BZ4 and BZ24/BZ19 recognized natural MAR and rhesus (RH) CRP, and only the recombinant MAR CRP. Additionally, the sensitivity of some mAb pairs such as BZ22/BZ24 was strong enough to require plasma samples to be diluted 2000 times to fall within the linear part of a standard curve. Thus, we were able to produce mAbs against MAR CRP that can be combined in different pairs depending on the species specificity of the ELISA assay.

**Figure 3.**
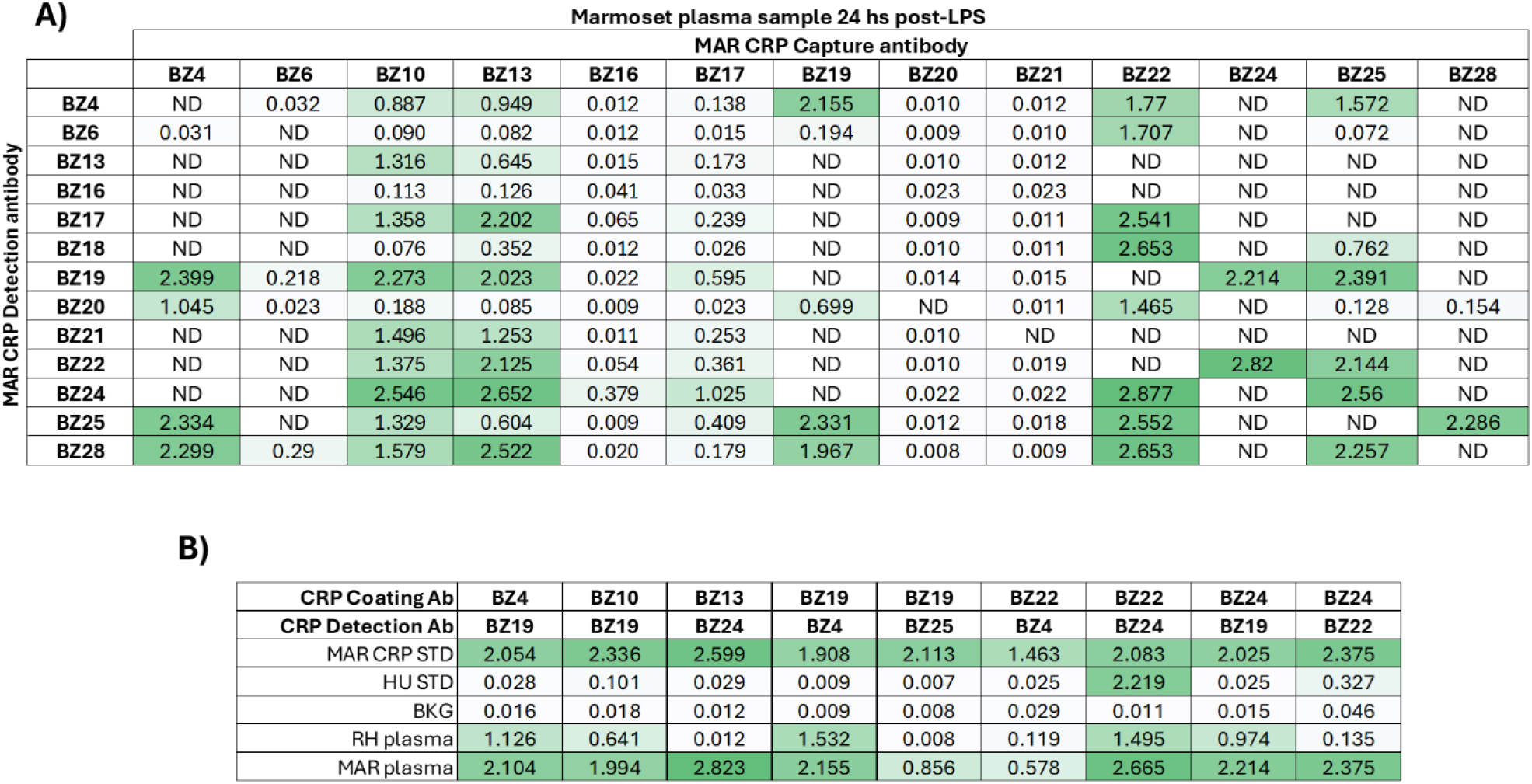
Screening of CRP-reactive mAbs using MAR-derived samples. A) Indirect ELISAs were set up with different capture/detection mAb combinations, and a 1:100 dilution of MAR plasma 24 hs post-LPS inoculation. B) Indirect ELISA with selected mAb pairs, using recombinant human or MAR standards or diluted (1:500) rhesus or MAR plasma. BKG: background signal. Values are optical densities measured at 450nm. ND: not done

MAR IL-4: A total of 10 mAbs were selected after screening using ELISA with recombinant MAR IL-4. These clones, and 2 additional ones that react with human and rhesus macaque IL-4 (IL-4I and IL-4II), were tested directly with MAR-derived samples. The antibodies described before were prepared in pure and biotinylated forms, and ELISPOT assays were performed to determine reactivity with MAR samples since IL-4 is generally released at low levels and consumed by cells in culture (Ewen and Baca-Estrada, 2001) (**Figure 4A**). Surprisingly, only the mAbs BR1, BR29, and IL-4(II) produced signals, with BR1 and BR29 targeting the same MAR IL-4 epitope (**Figure 2**). Based on the outcome of the first ELISPOT assay, antibodies BR29 and IL-4(II) were selected for further testing. Another ELISPOT assay was designed with PBMCs from three different marmosets, with or without stimulation with PHA, in which BR29 was used as capture Ab and IL-4(II) as detection Ab, or IL-4(II) as capture and BR29 as detection (**Figure 4B**). The results of this assay showed that either combination was able to detect IL-4-producing cells in the stimulated wells, but the combination BR29 capture with IL-4(II) detection produced clearer backgrounds.

**Figure 4.**
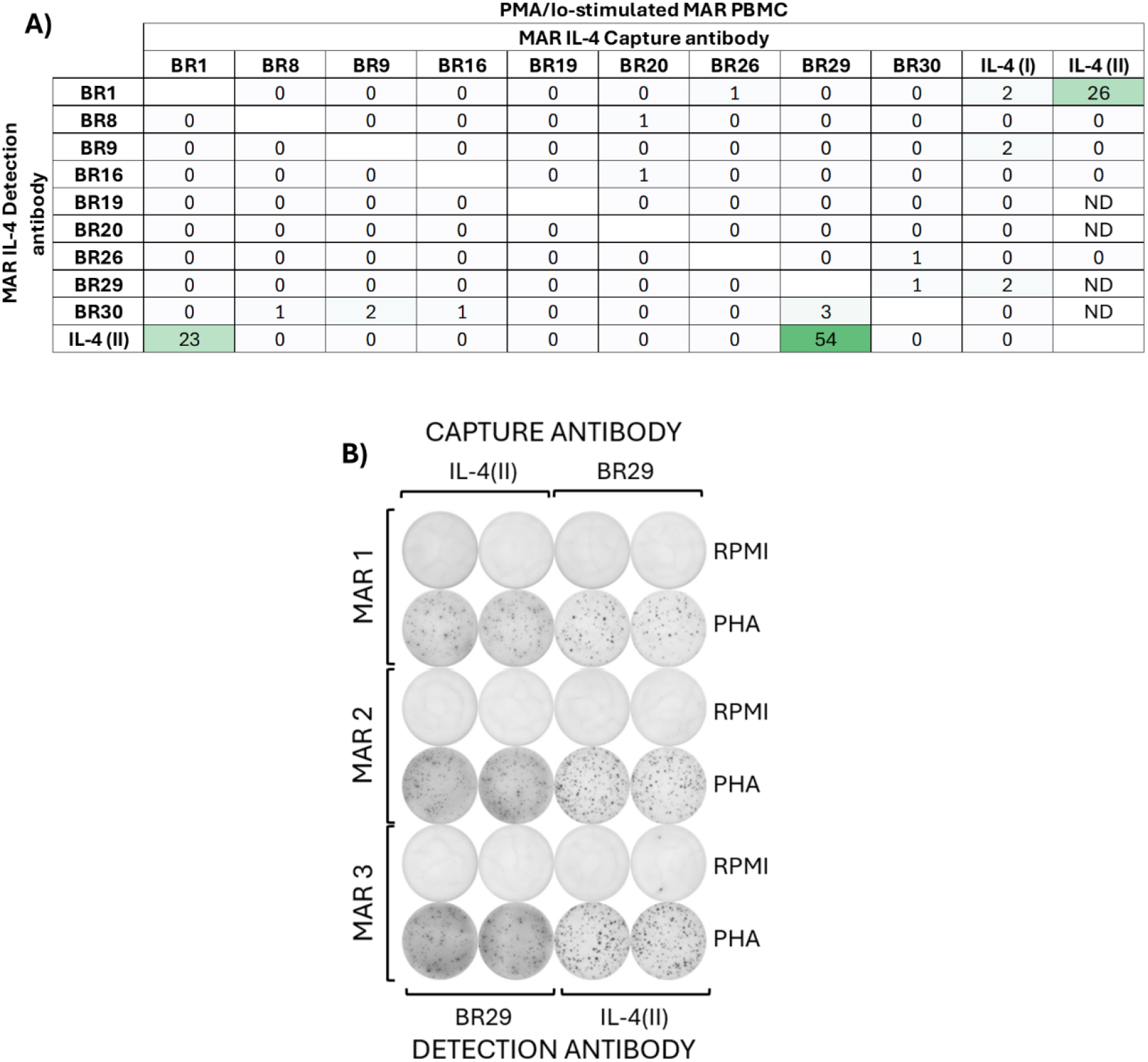
Screening of IL-4-reactive mAbs using MAR-derived samples. A) An ESLISPOT was set up with different capture/detection mAb combinations, and 5×10^3^ MAR PBMC/well stimulated with PMA/ionomycin for 48 hs. Spots were counted with an automated spot reader (CTL). B) The ELISPOT was repeated for BR29 and IL4(II) mAbs with 5×10^4^ PBMC/well from three different marmosets, with or without PMA/ionomycin stimulation for 48 hs. ND: not done.

MAR-IL-6: A total of nine purified and biotinylated mAbs, all reactive with recombinant MAR IL-6, were tested with natural MAR-derived samples; specifically, with supernatants of MAR PBMC stimulated with LPS, a TLR-4 ligand that induces expression of IL-6-in mammalian cells (Chow et al., 1999). There were only two mAb pairs that produced a significant signal, BU6/BU20 and BU7/BU13 (**Figure 5A**). These antibodies did not recognize recombinant human IL-6, or IL-6 produced by rhesus macaque and baboon PBMC (data not shown). BU6 and BU7 targeted the same epitope, and BU13 and BU20 targeted also the same epitope, but different from the one targeted by BU6 and BU7; however, the alternative combinations BU6/BU13 or BU7/BU20 failed to produce a detectable signal. We covalently bound BU6 or BU7 to LMX magnetic beads and performed assays with diluted supernatants of MAR PBMC unstimulated or stimulated with LPS. The highest fluorescence values were obtained with the mAb pair BU6/BU20, followed by BU7/BU13 (**Figure 5B**). Thus, we confirmed the identification of two antibody pairs that recognize both recombinant and natural MAR IL-6.

**Figure 5.**
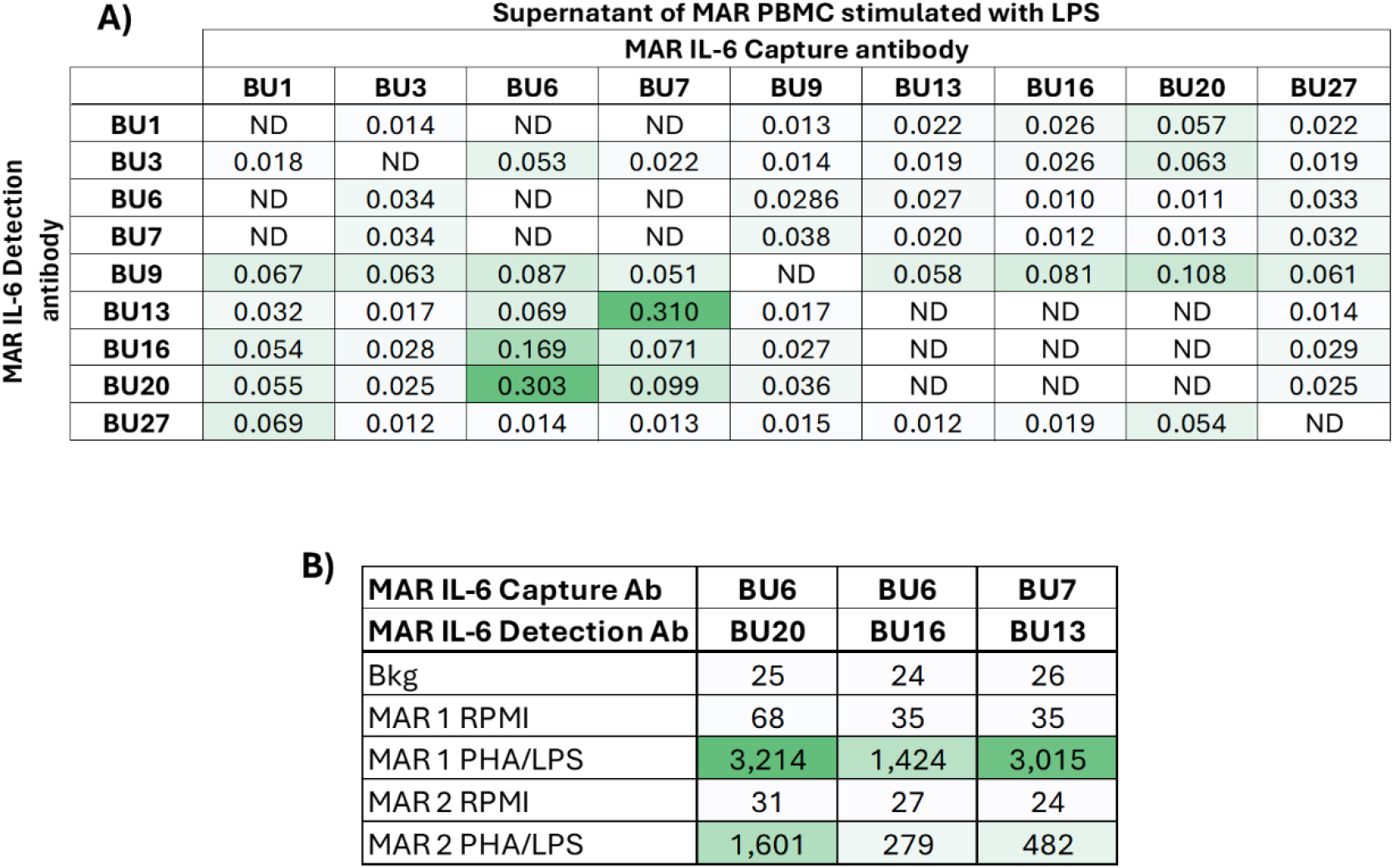
Screening of IL-6-reactive mAbs using MAR-derived samples. A) Indirect ELISAs were set up with different capture/detection mAb combinations, and a 1:10 dilution of supernatant of MAR PBMC 24 hs post-LPS stimulation. Values are optical densities measured at 450nm. ND: not done. B) Luminex (LMX) assay with selected mAb pairs, using 1:10 dilutions of MAR PBMC 24 hs post-LPS and PHA stimulation. Bkg: background signal. Values are median fluorescent intensities (MFI).

MAR IL-10: Antibody pairs that recognized two different epitopes of recombinant MAR IL-10 were identified (**Figure 6A**). Clone AD35 was covalently bound to LMX magnetic beads and combined with biotinylated AD32 to test for IL-10 expression from PHA-stimulated MAR PBMC, as well as similar archived samples from human, rhesus macaque, baboon, cynomolgus macaque, and African green monkey origin (Giavedoni, 2005). The combination AD35/AD32 produced signals from all the species tested (**Figure 6B**), as well as against recombinant human IL-10.

**Figure 6.**
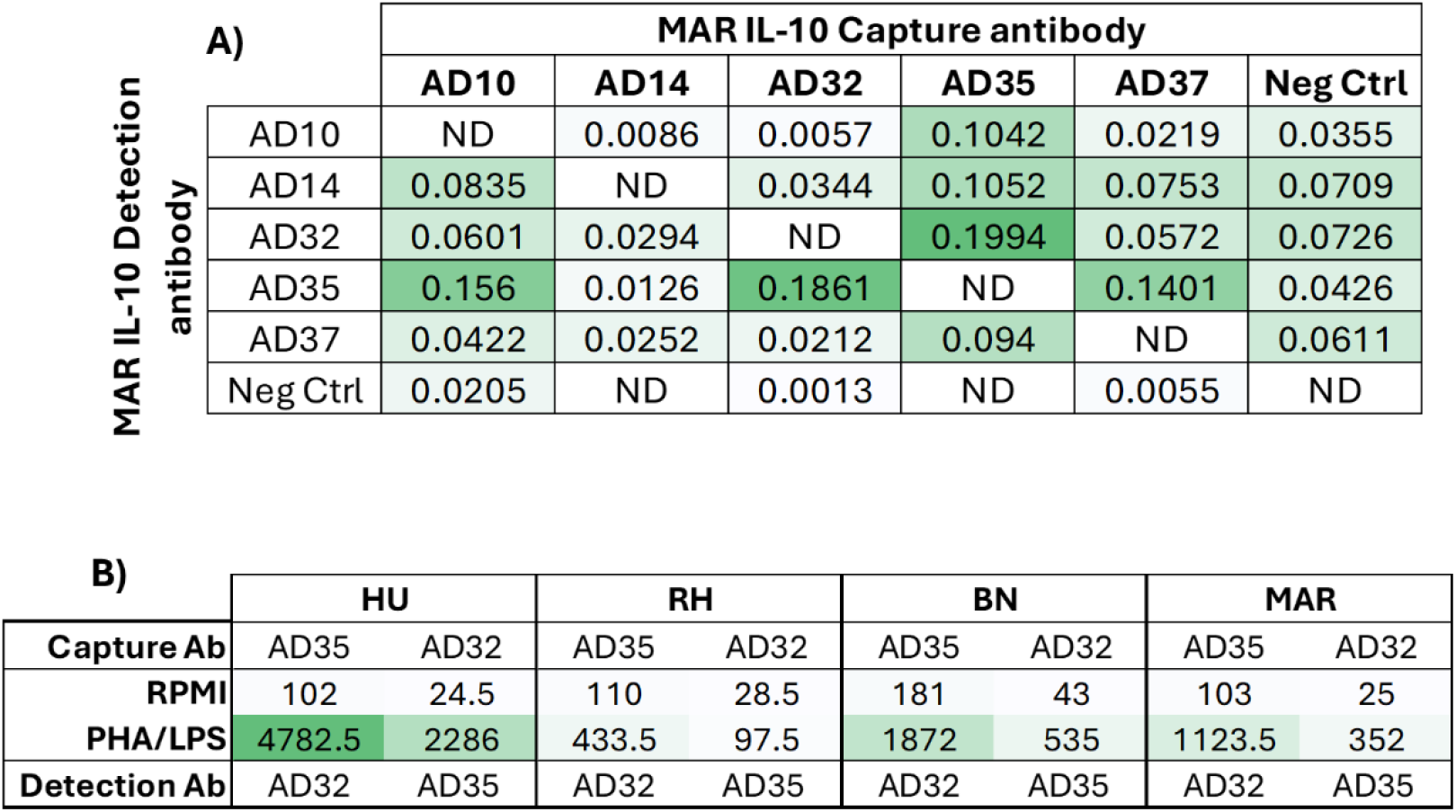
Screening of IL-10-reactive mAbs using MAR-derived samples. A) Epitope binning using Biolayer interferometry, soluble recombinant MAR IL-10 and different capture/detection combinations. ND: not done. B) Luminex (LMX) assay with AD32 and AD35 mAb pairs, using 1:10 dilutions of archived samples of human and NHP PBMC supernatants collected 24 hs post-LPS and PHA stimulation. HU: human; RH: rhesus macaque; BN: baboon; MAR: marmoset. Values are median fluorescent intensities (MFI).

MAR IP-10: A total of 14 mAbs, reactive against recombinant MAR IP-10, were processed and produced as pure and biotinylated forms, and ELISA assays were performed using all possible combinations; supernatants of MAR PBMCs stimulated with SEB were used for detection of working pairs (**Figure 7A**). The results showed that mAbs CD5, CD6, and CD7, which bind to different epitopes, were able to capture native MAR IP-10, and the combinations CD5/CD7, CD6/CD11, CD6/CD13, and CD7/CD5, detected the presence of MAR IP-10. MAbs CD6, CD7 and CD11 were covalently attached to LMX beads and a LMX assay was also run using different mAbs for detection (**Figure 7B**). The strongest MAR IP-10 signal originated when using CD11 as capture antibody and biotin-CD6 as detection antibody, which was unexpected considering that the CD11/CD6 combination had weak signals in the ELISA assay.

**Figure 7.**
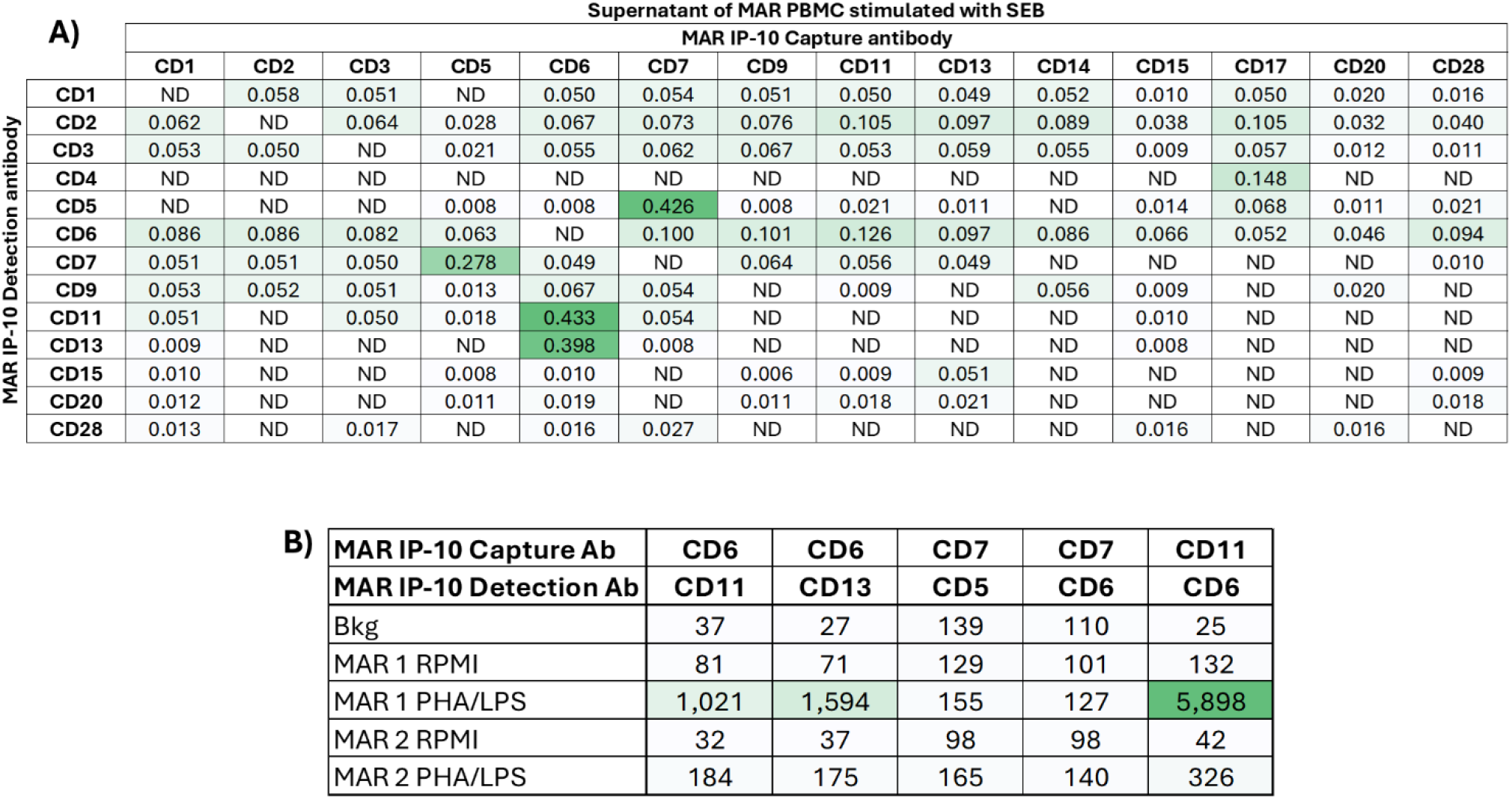
Screening of IP-10-reactive mAbs using MAR-derived samples. A) Indirect ELISAs was set up with different capture/detection mAb combinations, and a 1:10 dilution of supernatant of MAR PBMC 24 hs post-SEB stimulation. Values are optical densities measured at 450nm. ND: not done. B) Luminex (LMX) assay with selected mAb pairs, using 1:10 dilutions of MAR PBMC 24 hs post-LPS and PHA stimulation. Bkg: background signal. Values are median fluorescent intensities (MFI).

### Validation of selected mAb pairs

The validation of the anti-MAR IL-4 (BR29/IL-4 II) was demonstrated before, using an ELISPOT assay (**Figure 4B**). To validate the selected mAb pairs for MAR IL-6 (BU6/BU20), IL-10 (AD35/AD32), and IP-10 (CD11/CD6), we first selected supernatants of 20 MAR PBMC stimulated with a variety of stimulants that activate cells through different mechanisms. We tested PBMC from both female (10) and male (10) adult marmosets. We evaluated diluted supernatants collected at 24 hs post-stimulation, using a LMX assay that also included anti-human cytokine reagents that recognized MAR IFN-γ (MT-123L/7-B6-1) and TNF-α (MT21A8/MT15B15). The standard used for these assays included recombinant MAR IL-6 and IP-10, and recombinant human IFN-γ, IL-10, and TNF-α (**Figure 8**). The outcome of this LMX assay showed that PMA/ionomycin was the strongest inducer of IFN-γ and TNF-α, while LPS induced the highest levels of IL-6, and SEB the most IP-10; PMA/ionomycin and LPS increased the production of IL-10, but by a relatively small factor. There were no statistically significant differences in the levels of cytokines produced by male or female MAR PBMC (t-tests were performed for each cytokine and for each stimulant).

**Figure 8.**
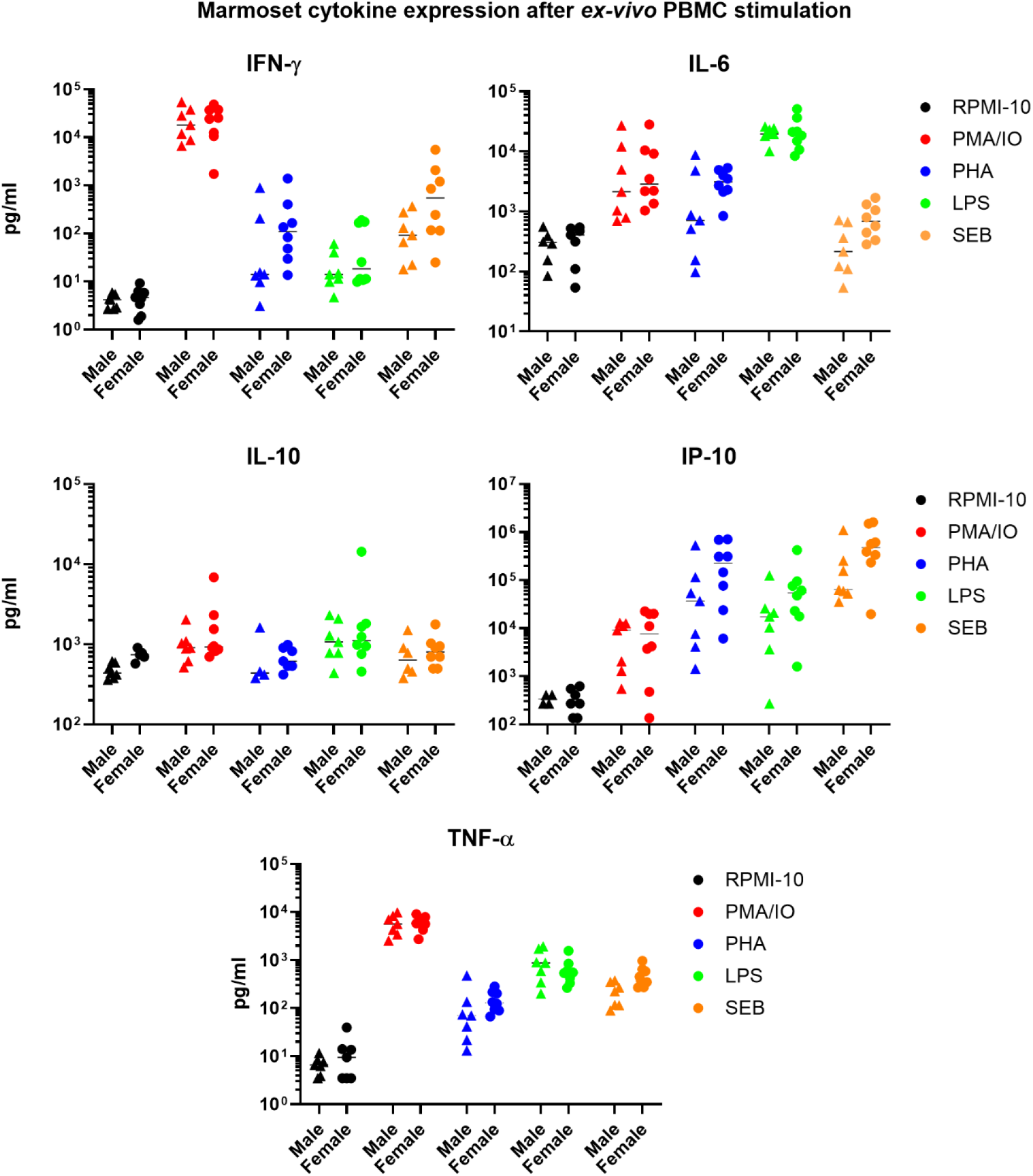
Validation of selected anti-MAR biomarker mAbs (capture mAb/detection mAb) with supernatant of stimulated MAR PBMC. A LMX assay was set up for detection of IFN-γ (MT123L/7-B6-1), IL-6 (BU6/BU20), IL-10, (AD35/AD32), IP-10 (CD11/CD6), and TNF-α (MT21A8/MT15B15) in the supernatant of MAR PBMC at 24 hs post-stimulation. Horizontal bars represent the median concentration for each group. T-tests were performed for male vs. female groups for each biomarker and each condition.

A second validation assay was performed by measuring the same five cytokines in the plasma of six MAR given a low-dose LPS intravenous inoculation. Plasma samples collected at different time-points were analyzed by the same LMX assay (**Figure 9**). The mAb pairs identified consistent changes for all animals, but different kinetics of systemic upregulation, depending on the cytokine being analyzed. For IFN-γ, there were no significant changes in plasma concentration until about 2 hs post-LPS injection, when a steady increase was noticed, which peaked at about 4-6 hs post-LPS injection and returned to baseline at 24 hs. In the case of IL-6, an increase in concentration was already observed 30 min post-LPS injection, with concentration peaking at 1-2 hs and declining afterwards, although half of the animals had IL-6 levels above baseline by 24 hs. For IL-10, like for IL-6, there were detectable increases by 30 min post-LPS injection; however, the peak of expression occurred at 1 h, and baseline levels were reached by 6 hs post-LPS injection. For IP-10, there was a 30 min delay in upregulation, but the peak of concentration lasted from 2 hs to more than 6 hs post-LPS injection, descending to baseline levels by 24 hs. Finally, for TNF-α the peak of concentration lasted from 1 h to 2 hs post-LPS injection, and it returned to baseline by 6 hs.

**Figure 9.**
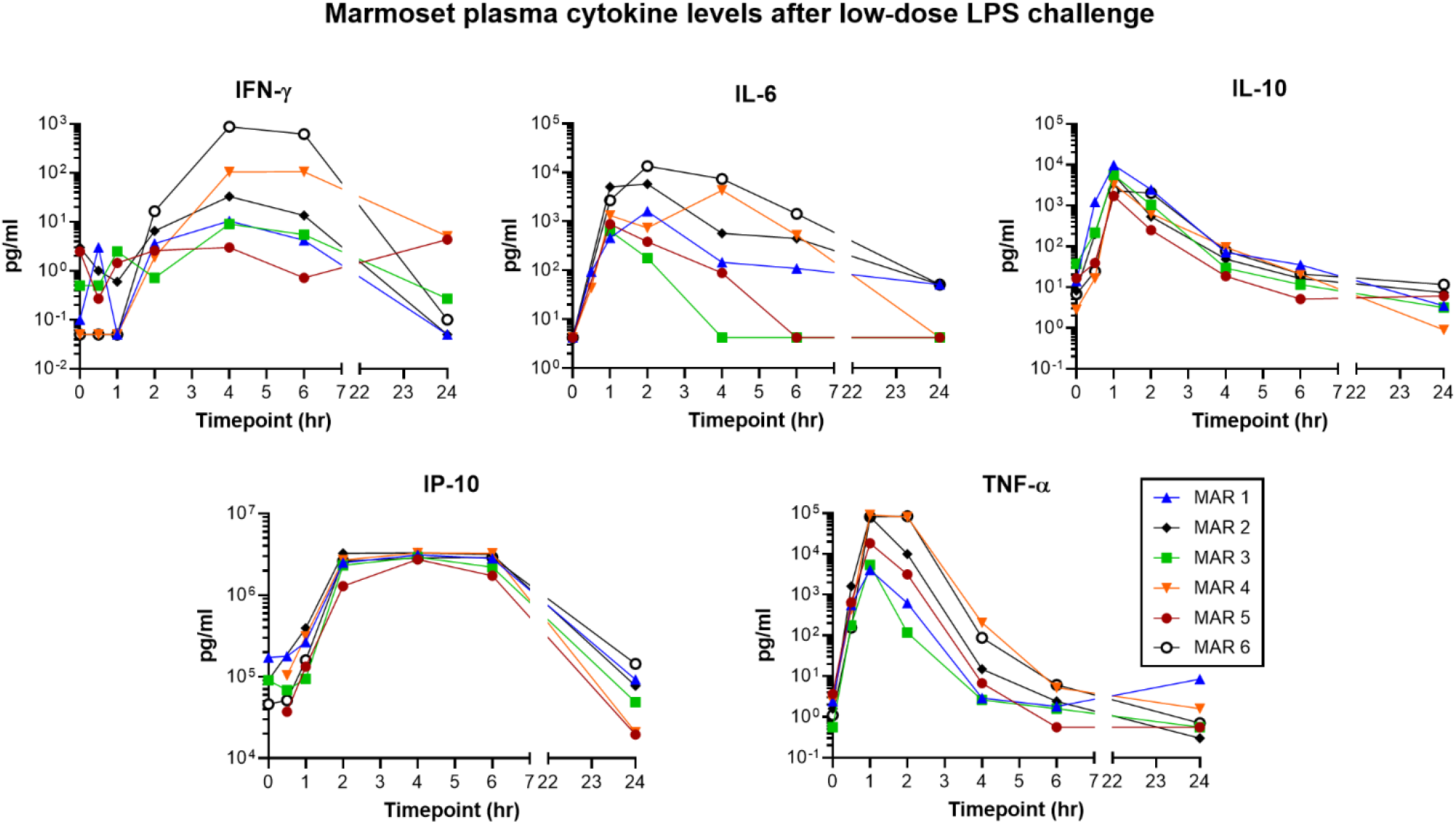
Validation of selected anti-MAR biomarker mAbs (capture mAb/detection mAb) with plasma of MAR undergoing low-dose LPS intravenous challenge. A LMX assay was set up for detection of IFN-γ (MT123L/7-B6-1), IL-6 (BU6/BU20), IL-10 (AD35/AD32), IP-10 (CD11/CD6), and TNF-α (MT21A8/MT15B15).

Finally, for CRP we compared plasma concentrations in five MAR before LPS challenge and at 24 hs post inoculation (**Figure 10**). The levels of CRP detected in the MAR plasma at 24 hs showed a statistically significant increase after LPS exposure (p<0.0001, two-way Anova), suggesting the proper specificity of the assay. Interestingly, MAR 3, who had the lowest increase in CRP, was the same animal that most rapidly had IL-6 levels returning to baseline values (4 hs post-LPS challenge).

**Figure 10.**
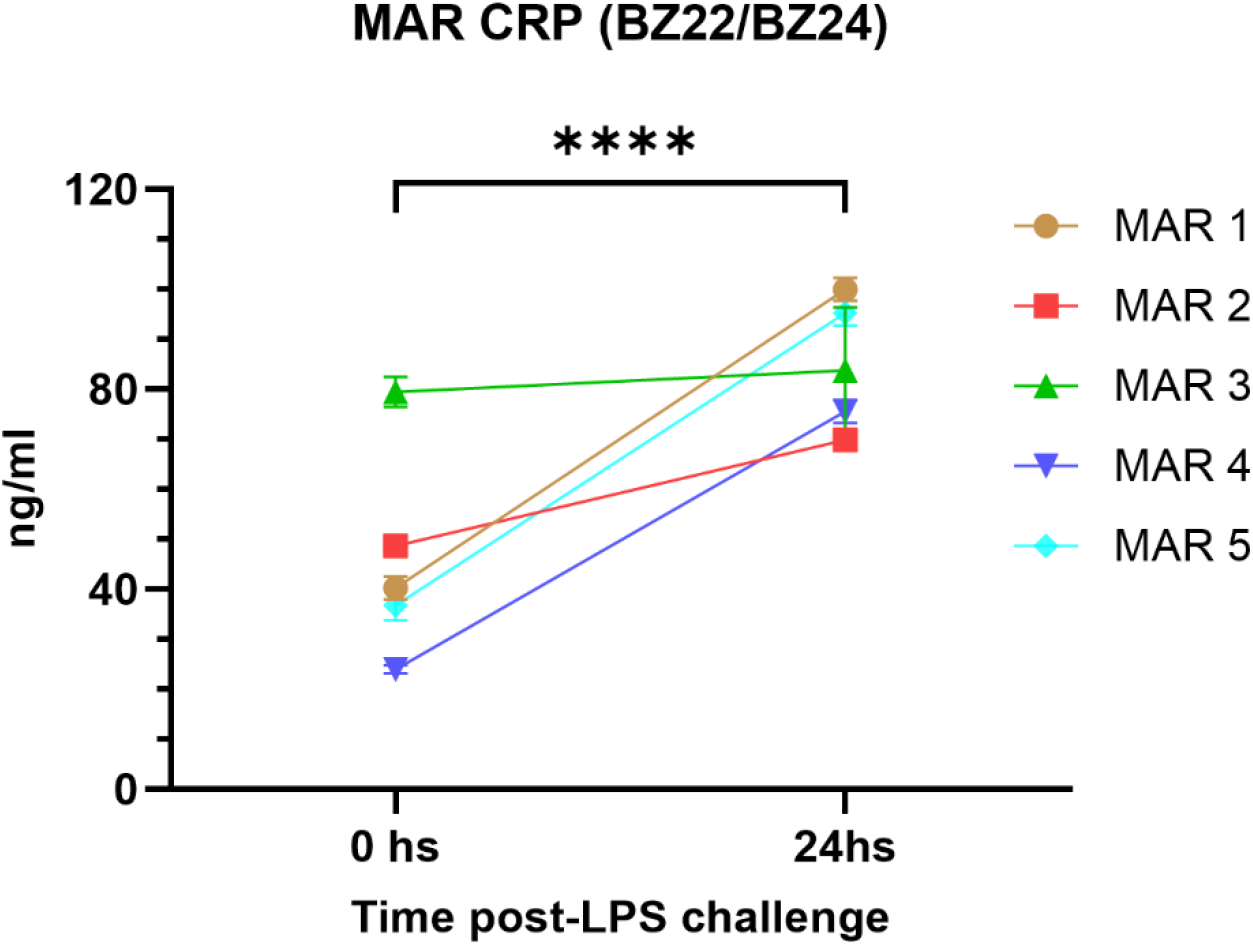
Validation of a selected anti-MAR CRP mAb pair with MAR plasma. An ELISA was set up with the BZ22 capture antibody and biotinylated BZ24 detection antibody. Plasma from five MAR challenged with a low-dose LPS intravenous challenge, at 0 and 24 h, was diluted 1:2000 for detection. Human CRP (RCD Systems) was used for the standard curve. A two-way Anova analysis was performed to determine significant changes after LPS treatment (****: p<0.0001).

In summary, we were able to identify working mAb pairs in different immunoassays for all five biomarkers of inflammation, which detected increases in MAR-derived samples after different types of stimulation.

## Discussion

Inflammation is a common process involved in aging, infectious disease, autoimmune diseases such as multiple sclerosis (MS), and age-related neurodegenerative disorders such as Alzheimer’s disease (AD) and Parkinson’s disease (PD), all of which are being modeled in the MAR. Most of the immunological assays currently available to identify biomarkers of inflammation in NHP species are based on the use of mAbs developed against human antigens. While several markers have been validated for use in MAR models (Giavedoni, 2005, Mustoe et al., 2026, Peters et al., 2023, Singh et al., 2021, Ross et al., 2019, Seferovic et al., 2018, Manickam et al., 2017, Hoglind et al., 2017, Chiu et al., 2017, Manickam et al., 2016), there were a large number of markers that failed. The goal of this study was to improve the translational value of MAR by generating mAbs and developing immunoassays to identify important biomarkers of inflammation, for which there is currently scarcity of reagents for this species. Mouse immunization involved the fabrication of several variants of the MAR biomarkers. While these recombinant proteins were used to identify the capacity of the newly developed mAbs to bind the same or similar antigen variants used for immunization, the ultimate test was whether the same antibodies would recognize the native molecule in biological samples from MAR, which were produced both by in-vivo and ex-vivo approaches.

CRP was first identified in 1930 as a substance in the serum of patients with acute inflammation that reacted with the “C” carbohydrate antibody of the capsule of pneumococcus (Tillett and Francis, 1930). CRP is a pentameric protein synthesized by the liver and is a protein primarily induced by the action of IL-6 on the gene responsible for transcription of CRP during the acute phase of an inflammatory/infectious process (Nehring and Patel, 2019, Nagasawa, 2026). The large number of different epitopes that we identified for MAR CRP in this study may be due to the polymeric nature of this protein. Interestingly, we observed that some mAb pairs only recognized the MAR CRP molecule, while others recognized more conserved epitopes that are present in MAR proteins from several species; however, the affinity of the new mAbs with these conserved epitopes may be different for different species and was not evaluated in this study.

IL-4 has an important role in regulating antibody production, hematopoiesis and inflammation, and the development of effector Th2-cell responses (Brown and Hural, 2017). We selected an ELISPOT format that can identify IL-4-producing cells as soon as the cytokine is released into the supernatant. In our study, the recombinant MAR IL-4 variants that we produced induced numerous antibodies that reacted with these proteins, but just two mAbs (BR1 and BR29) that reacted with the same epitope of the native form of MAR IL-4. Fortunately, we found an anti-human IL-4 antibody (IL-4II) that recognized MAR IL-4 and could be paired with BR29 to produce a sensitive ELISPOT assay.

IL-6 is an important mediator of fever and of the acute phase response, capable of crossing the blood-brain barrier and initiating synthesis of PGE_2_ in the hypothalamus, thereby changing the body’s temperature set-point (Clark, 1989). IL-6 is secreted by macrophages in response to pathogen-associated molecular patterns (PAMPs) antigens, small molecules, monoclonal antibodies and immune checkpoint inhibitors (Unver and McAllister, 2018). IL-6 is an important biomarker used in infectious and autoimmune diseases, cancer, neurodegenerative and cardiovascular diseases, and aging (Erol, 2007). As described above for IL-4, screening of the large number of mAbs with anti-recombinant MAR IL-6 binding activity reduced the number of mAb pairs that recognized the native MAR IL-6 form to just three pairs. Another interesting outcome was that the same mAb pairs produced different affinities for natural MAR IL-6 depending on whether they were used in an ELISA or an LMX assay. For example, BU7/BU13 generated a slightly stronger signal than BU6/BU20 in ELISA format, but BU6/BU20 produced the strongest signal in the LMX format and was selected for all the analyses that used that assay. This difference in signal may reflect the conformational changes that mAbs (BU6 and BU7) experience when immobilized to a solid support by electrostatic forces (ELISA), compared to covalent bonds binding antibodies to carboxylated beads (LMX).

IL-10 is a cytokine expressed mainly by monocytes with pleiotropic effects in immunoregulation and inflammation. IL-10 was initially reported to suppress cytokine secretion, antigen presentation and CD4+ T cell activation, but further investigation has shown that IL-10 predominantly inhibits lipopolysaccharide (LPS) and bacterial product mediated induction of the pro-inflammatory cytokines TNF-α, IL-1β, IL-12, and IFN-γ secretion from Toll-Like Receptor (TLR) triggered myeloid lineage cells (Ouyang and O’Garra, 2019). In this study, IL-10 production by ex vivo stimulated MAR PBMC was rather modest, but this could have reflected a different kinetic expression for IL-10, since supernatants were collected at 24 hs post-stimulation. Still, LPS was one of the stimulants that induced a higher IL-10 expression. In an in vivo setting, LPS challenge induced a rapid IL-10 systemic response that paralleled the LPS-induced expression of IL-6 and TNF-α. The mAb pairs produced in this study (AD35/AD32) recognized conserved IL-10 epitopes that are present in the human, as well as MAR and some Old-World Monkey species homologues.

IP-10 (CXCL10) is a chemokine secreted from cells stimulated with type I and II interferons (IFNs) and lipopolysaccharide (LPS) that acts as chemoattractant for activated T cells into sites of tissue inflammation (Dufour et al., 2002). IP-10 has been recently identified as a sensitive biomarker for inflammatory conditions including among others, development and treatment of tuberculosis (Ruhwald et al., 2009, Goosen et al., 2015), acute respiratory infections (Hayney et al., 2017), Kawasaki disease (Ko et al., 2015), viral infections (Quint et al., 2010), and cystic fibrosis exacerbation (Solomon et al., 2013). Binding of the generated mAbs to native MAR IP-10 was seen for a minority of mAb pairs, and signal strength also changed for the same mAb pairs depending on the assay being run (ELISA vs LMX). The selected CD11/CD6 mAb pair gave a very strong signal in the LMX format and was used for all the validation assays.

The ligand binding assays (LBA), such as ELISA and LMX, are the archetypical quantitative assays for biomarkers. However, as most biomarkers are endogenous substances and already present in samples, an analyte-free matrix to utilize during validation studies is more difficult to obtain. Access to a fully characterized form of biomarker to act as a calibration standard is also limited. Thus, most available biomarkers LBA fall into the category of relative quantitation (Lee et al., 2005). The FDA and NIH Joint Leadership Council developed the BEST (Biomarkers, EndpointS, and other Tools) Resource (2016), which defined validation as “a process to establish that the performance of a test, tool, or instrument is acceptable for its intended purpose”. Validation thus informs essentially all potential uses of biomarkers, establishing whether biomarkers (and the tests used to assess them) are “fit-for-purpose” and appropriate for specific contexts of use in product development and risk assessment. This definition, the lack of “gold standard” for most protein biomarkers (Neely, 2018), the different biological relevance of the proteins that we selected as targets in this study, and the varied number of research interests with the MAR, indicates that there is not a single process for validating a specific biomarker that will satisfy all of them. Ultimately, fit-for-purpose requires an assessment of the technical ability of the assay to deliver against the predefined purpose (Cummings et al., 2010). Thus, we selected as fit-for-purpose validation feature for our mAbs the production of a specific signal in stimulated MAR samples that is significantly higher than the signal detected in unstimulated samples, and the selected mAbs that we report in this study fulfil this requirement.

## Conclusion

We successfully produced and identified mAb pairs that detect biomarkers of inflammation in MAR biological samples. Access to these reagents for the scientific community is available at https://trinity.edu/nwmimmunoreagents. The most appropriate immunoassay varies depending on the target: ELISPOT for MAR IL-4, ELISA for MAR CRP, and LMX for MAR IL-6, IL-10 and IP-10. With the addition of mAb pairs for IFN-γ and TNF-α, the LMX reagents allow for the detection of five MAR biomarkers in very small plasma volumes, which is very convenient considering the limited blood volume that can be drawn from these small NHPs. These reagents are expected to improve the translational value of MAR-based biomedical models of inflammatory conditions.

## Acknowledgments

The authors would like to acknowledge veterinarians, veterinary technicians, and personnel at the Karolinska Institute and the SNPRC for their continued support.

## Authorship confirmation/contribution statement

E.L.: Investigation, Analysis, Writing – review & editing; D.H.: Investigation, Analysis, Writing – review & editing; B.T.: Investigation, Analysis; J.C.: Resources, Investigation; V.H.: Analysis, Writing – review & editing; J.K.: Investigation, Analysis; C.R.: Resources, Writing – review & editing; N.A.: Conceptualization, Funding Acquisition, Supervision, Methodology; B.M.: Conceptualization, Resources, Supervision, Methodology, Writing – review & editing; L.D.G.: Conceptualization, Resources, Analysis, Funding Acquisition, Supervision, Writing – original draft, Writing – review & editing, Project Administration.

## Reagent Availability Statement

The monoclonal antibodies and immunoassays described here are available to the scientific community as indicated at https://trinity.edu/nwmimmunoreagents. The recombinant antigen variants listed in Supplemental Table 1 and Table 2 are available from Mabtech AB to qualified investigators under a material transfer agreement. Mabtech AB will consider confidential disclosure of the proprietary immunogenicity-enhancing elements to the editor on request.

## Animal Studies Statement

All the experiments involving mice were performed at the Karolinska Institute, Solna, Sweden, according to the guidelines of the Swedish Ethical Committee for Animal Protection. Marmoset studies were performed at the Southwest National Primate Research Center following protocols approved by Institutional Animal Care and Use committee of the Texas Biomedical Research Institute, San Antonio, TX, USA.

---

**Supplemental Table 1.**
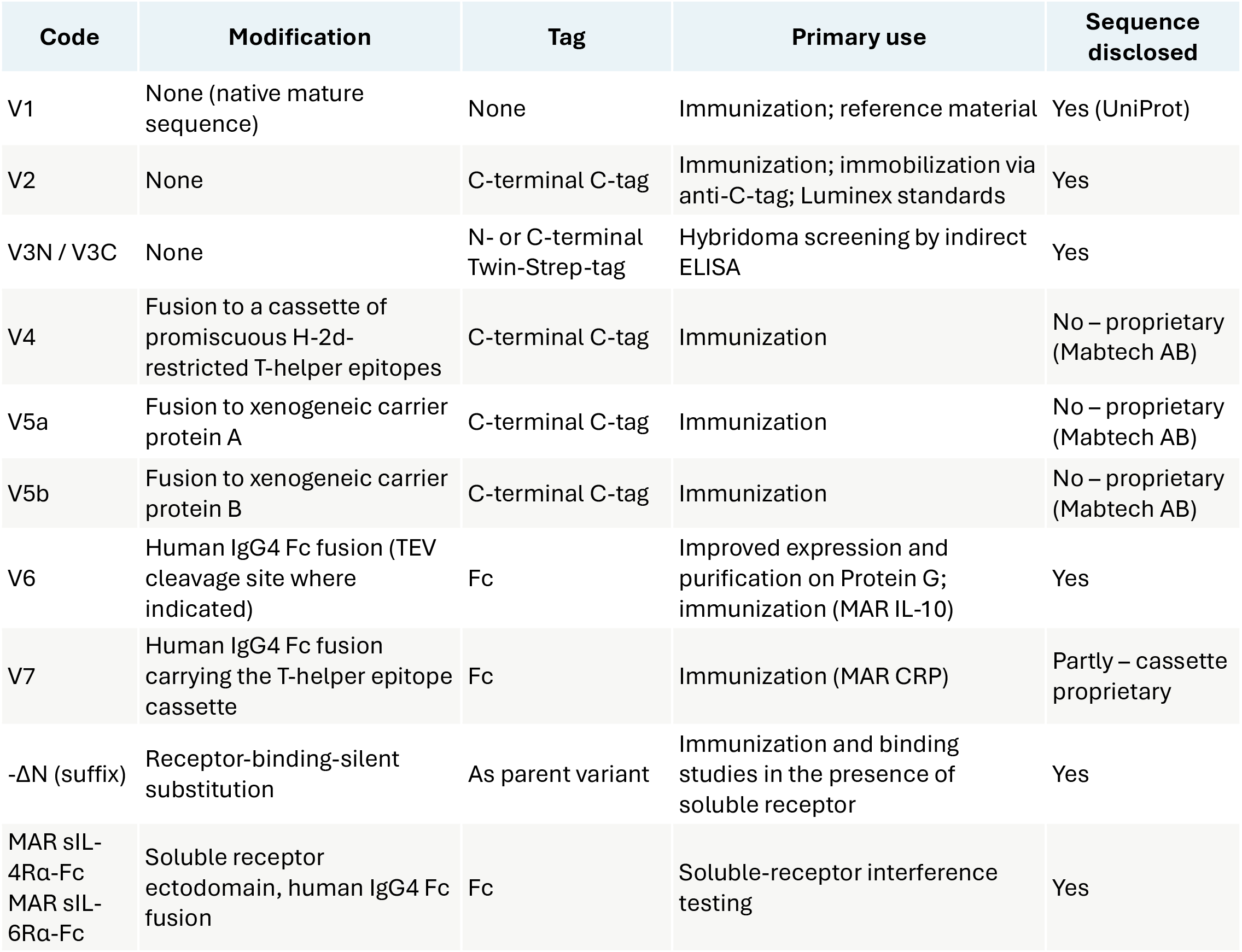
Recombinant marmoset antigen variants generated in this study, and the codes used to designate them throughout the text, figures and tables. All variants were expressed in HEK-293 cells. Variants V4, V5a, V5b and V7 incorporate proprietary immunogenicity-enhancing elements of Mabtech AB whose identity and sequence are not disclosed (see Author Disclosure Statement); the general strategies of T-helper epitope cassette fusion and of fusion to heterologous immunostimulatory proteins are described elsewhere (Alexander et al., 1994; Hung et al., 2007; McCormick et al., 2001; Nimal et al., 2005). Expression plasmids for all variants are listed in Supplementary Table 1.

**Supplemental Table 2.** Recombinant MAR antigen constructs and expression plasmids generated for this study. Constructs are identified by the variant codes defined in Supplementary Table 1. The identity and sequence of the proprietary immunogenicity-enhancing elements carried by variants V4, V5a, V5b and V7 are not disclosed (see Author Disclosure Statement).

| Antigen | Variant | Modification | Tag | Expression plasmid | Calc. MW (kDa) |
| --- | --- | --- | --- | --- | --- |
| IL-10 | V6 | Human IgG4 Fc fusion, TEV site | Fc | pcDNA 3.1 Zeo- | 47,9 |
| IL-10 | V1 | None | None | pcDNA 3.1 Zeo- | 18,7 |
| IL-10 | V3N | None | N-terminal Twin-Strep | pcDNA 3.1 Zeo- | 21,8 |
| IL-10 | V3C | None | C-terminal Twin-Strep | pcDNA 3.1 Zeo- | 21,6 |
| IL-4 | V2 | None | C-tag | pcDNA 3.1 Zeo- | 15,2 |
| IL-4 | V2-ΔN | Receptor-binding-silent | C-tag | pcDNA 3.1 Zeo- | 14,8 |
| IL-4 | V4 | T-helper epitope cassette (proprietary) | C-tag | pcDNA 3.1 Zeo- | 19,6 |
| IL-4 | V5a | Xenogeneic carrier protein A (proprietary) | C-tag | pcDNA 3.1 Zeo- | 31,2 |
| IL-4 | sIL-4Rα-Fc | Soluble receptor ectodomain, IgG4 Fc fusion | Fc | pcDNA 3.1 Zeo- | 48,2 |
| IL-4 | V3N | None | N-terminal Twin-Strep | pcDNA 3.1 Zeo- | 18,0 |
| IL-4 | V3C | None | C-terminal Twin-Strep | pcDNA 3.1 Zeo- | 17,8 |
| IL-4 | V1 | None | None | BM.3 | 14,8 |
| IL-6 | V2 | None | C-tag | pcDNA 3.1 Zeo- | 21,3 |
| IL-6 | V2-ΔN | Receptor-binding-silent | C-tag | pcDNA 3.1 Zeo- | 20,9 |
| IL-6 | V4 | T-helper epitope cassette (proprietary) | C-tag | pcDNA 3.1 Zeo- | 25,7 |
| IL-6 | V5a | Xenogeneic carrier protein A (proprietary) | C-tag | pcDNA 3.1 Zeo- | 37,3 |
| IL-6 | sIL-6Rα-Fc | Soluble receptor ectodomain, IgG4 Fc fusion | Fc | pcDNA 3.1 Zeo- | 64,8 |
| IL-6 | V3N | None | N-terminal Twin-Strep | pcDNA 3.1 Zeo- | 24,1 |
| IL-6 | V3C | None | C-terminal Twin-Strep | pcDNA 3.1 Zeo- | 23,9 |
| IL-6 | V1 | None | None | BM.3 | 20,9 |
| IP-10 | V2 | None | C-tag | pcDNA 3.1 Zeo- | 9,2 |
| IP-10 | V2-ΔN | Receptor-binding-silent | C-tag | pcDNA 3.1 Zeo- | 8,7 |
| IP-10 | V4 | T-helper epitope cassette (proprietary) | C-tag | pcDNA 3.1 Zeo- | 13,5 |
| IP-10 | V5a | Xenogeneic carrier protein A (proprietary) | C-tag | pcDNA 3.1 Zeo- | 25,2 |
| IP-10 | V3N | None | N-terminal Twin-Strep | pcDNA 3.1 Zeo- | 11,9 |
| IP-10 | V3C | None | C-terminal Twin-Strep | pcDNA 3.1 Zeo- | 11,8 |
| IP-10 | V1 | None | None | BM.3 | 8,8 |
| CRP | V2 | None | C-tag | pcDNA 3.1 Zeo- | 23,2 |
| CRP | V4 | T-helper epitope cassette<br>(proprietary) | C-tag | pcDNA 3.1 Zeo- | 27,7 |
| CRP | V5a | Xenogeneic carrier protein A<br>(proprietary) | C-tag | pcDNA 3.1 Zeo- | 39,4 |
| CRP | V5b | Xenogeneic carrier protein B<br>(proprietary) | C-tag | pcDNA 3.1 Zeo- | 39,6 |
| CRP | V7 | Human IgG4 Fc fusion + T-helper<br>epitope cassette | Fc | pcDNA 3.1 Zeo- | 54,6 |
| CRP | V3N | None | N-terminal Twin-Strep | pcDNA 3.1 Zeo- | 26,2 |
| CRP | V3C | None | C-terminal Twin-Strep | pcDNA 3.1 Zeo- | 25,8 |

**Supplemental Figure 1.**
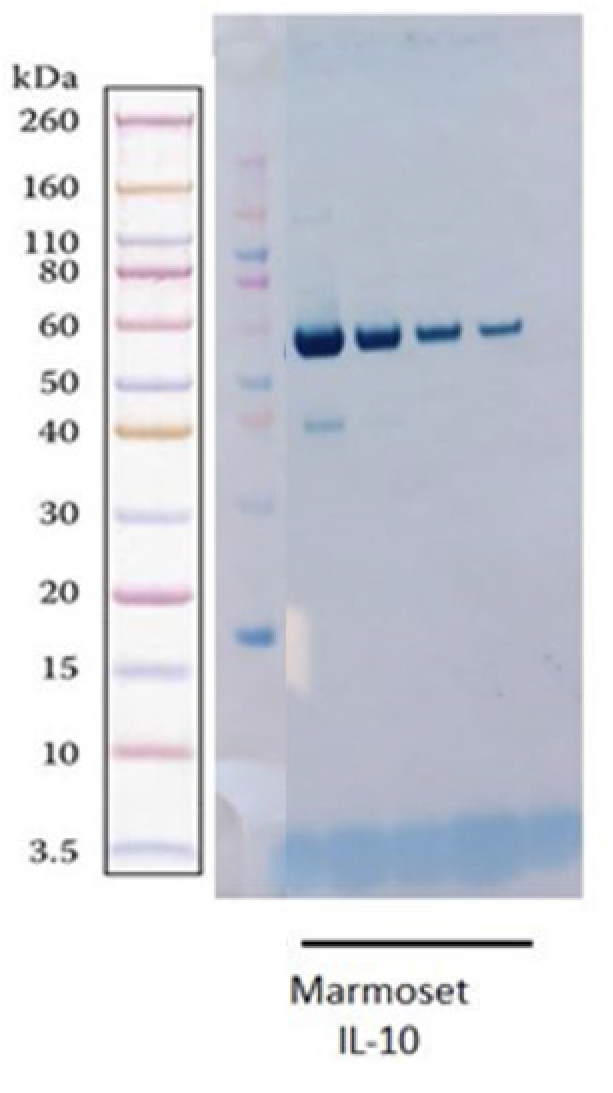
Expression and purification of a MAR IL-10 variant. MAR IL-10 was expressed in HEK293 cells as a fusion protein with the human IgG4 Fc fragment (variant V6, Table 1). The protein was purified using protein G and analyzed using SDS-PAGE and western blot with an anti-human IgG antibody. The lanes were loaded with different amounts of purified protein.

